# Risk assessment of zoonotic viruses in urban-adapted wildlife

**DOI:** 10.1101/2024.12.18.629064

**Authors:** Xuemin Wei, Hongfeng Li, Zheng Y.X. Huang, Shuo Li, Yuhao Wang, Jie Lan, Li Hu, Yang Li, Daniel J. Becker, Fuwen Wei, Yifei Xu

**Affiliations:** Department of Microbiology, School of Public Health, Cheeloo College of Medicine, Shandong University, Jinan, Shandong, China; Department of Zoology, School of Life Sciences, Nanjing Forestry University, Nanjing, Jiangsu, China; CAS Key Laboratory of Animal Ecology and Conservation Biology, Institute of Zoology, Chinese Academy of Sciences, Beijing, China; School of Biological Sciences, University of Oklahoma, Norman, OK, USA; Jiangxi Provincial Key Laboratory of Conservation Biology, College of Forestry, Jiangxi Agricultural University, Nanchang, Jiangxi, China

**Author notes:** Contributed equally (Xuemin Wei, Hongfeng Li, Zheng Y.X. Huang).

**Keywords:** urban adaptation, virus diversity, zoonotic potential, risk assessment, urbanization, spillover, spillback

## Abstract

The repeated emergence of pandemic viruses underscores the linkages between land-use change and wildlife disease, and urban-adapted wildlife are of special interest due to their close proximity to humans. However, viral diversity within urban-adapted species and their zoonotic potential remains largely unexplored. We compiled a large dataset on seven priority urban-adapted mammal species and their viruses covering over 115 countries from 1574 to 2023. These urban-adapted species host 286 virus species spanning 24 orders and 38 families, 14 of which are potentially high risk for human infection. Raccoons carried the most high-risk viruses, while raccoon dogs had increased viral positivity in urban habitats compared to raccoons, wild boars, and red foxes. Many viruses in urban-adapted species were phylogenetically related to those found in humans, and we also observed evidence of possible viral spillback. These results highlight zoonotic risks associated with urban-adapted species and suggest enhanced surveillance to mitigate future outbreaks.

## Introduction

Cities are not only home to the world’s human population but also host a diverse array of wildlife. Urban-adapted wildlife are those species that have adapted to living in cities and suburban areas^1,2^ With more animals affected by climate change amid an ever-expanding urban landscape, it is increasingly common to see wildlife sharing the same space with humans in urban habitats^3^.

Most human infectious diseases are caused by viruses originating from wildlife through cross-species transmission, and anthropogenic land-use change is one of the primary drivers of zoonotic spillover^4–6^. Urban-adapted wildlife are of special interest due to their close proximity and frequent interactions with humans^7,8^. The COVID-19 pandemic has further underscored the potential importance of urban-adapted wildlife to global health, given how wildlife in human interfaces have been linked to the origin of SARS-CoV-2^6,9,10^. Urban-adapted species may serve as donor or amplifier hosts for zoonotic viruses, and the dynamic interactions between these hosts, domestic animals, and humans increase the opportunities for virus spillover^11,12^. Conversely, human viruses could also spread to urban-adapted wildlife (i.e., spillback), opening pathways for the emergence of novel variants with unpredictable characteristics that could eventually be transmitted back to humans^13–15^. Therefore, urban-adapted species may play a neglected but crucial role in wildlife viral ecology and zoonotic risk assessment. A better understanding of zoonotic viruses in urban-adapted species is of central importance for mitigating and preventing future infectious disease outbreaks.

Previous research has identified a spectrum of viruses in urban-adapted wildlife, some of which may present high risks to humans^16–24^. Prior work also has found that urban-adapted mammal species carry more zoonotic pathogens than non-urban species^25,26^. While these studies have provided important insights into viral discovery and diversity, they commonly focus on a single virus or a small number of viruses of interest. Similarly, such studies often compare virus outcomes at either intra-specific (i.e., differences among rural and urban populations) or inter-specific (i.e., differences among urban-adapted and non-adapted species) scales of analysis. A thorough understanding of virus diversity and positivity within urban-adapted species is lacking. Moreover, little is known about virus sharing between urban-adapted species and humans at wildlife–human interface and the zoonotic potential of these shared viruses.

This knowledge gap is fundamental for better predicting epidemics and pandemics and for designing and implementing practical infection control strategies.

Here, we investigated the risk of zoonotic viruses in seven priority urban-adapted mammal species from three orders, including *Vulpes vulpes* (red fox), *Procyon lotor* (raccoon), *Nyctereutes procyonoides* (raccoon dog), and *Paguma larvata* (masked palm civet, hereafter palm civet) from the order Carnivora; *Erinaceus europaeus* (European hedgehog) and *Sorex araneus* (European shrew) from the order Eulipotyphla; and *Sus scrofa* (wild boar) from the order Artiodactyla. These seven priority urban-adapted species have shown rapid population growth and range expansion over the past decades^27–30^. In addition, these species have been linked with high-profile zoonotic viruses, including but not limited to highly pathogenic avian influenza (HPAI) H5N1 virus, severe acute respiratory syndrome (SARS)-related coronavirus, and severe fever with thrombocytopenia syndrome virus (SFTSV), raising concern about their potential threat to public health^31–33^. We compiled and then analyzed a large and novel dataset on these urban-adapted species and their associated viruses through literature review and analyses of public data, comprising more than 1,658,000 records from over 115 countries spanning 1574 to 2023. Our objectives were to unveil the diversity and global distribution pattern of viruses in these priority urban-adapted species, test how urbanization affects viral positivity within each species, and assess the potential risk of pathogenic viruses to humans. Our work highlights urban-adapted mammals that present risk of viral spillover to humans and informs risk-based allocations of research and surveillance.

## Methods

### Data collection

We collected data on seven priority urban-adapted species. The habitat preferences of these seven species were determined based on a published database of long-term, urban- adaptation status across mammals^2^. Data were compiled from multiple sources, including literature reviews, relevant websites, and public databases.

For the literature review, two reviewers independently and systematically searched electronic databases (PubMed and China National Knowledge Infrastructure) to extract data from research papers published before December, 2023. Following a systematic procedure^34^, publications containing the Latin and common names of these seven mammal species in the title and abstract were retrieved, including 34,394 English publications and 9,710 Chinese publications (Fig. S1). Reviews and duplicate publications were excluded. We retained 997 articles containing detailed information on the geographic distribution of these urban-adapted species and their associated viruses. From each publication, we extracted the title, authors, year of publication, research location (geographical coordinates or the most precise description), study period (the timeframe of sample collection), area type (urban areas, natural areas, or unknown), host species, sample type, organ distribution, detected virus species, detection methods, total number of samples, and number of positive samples. Areas corresponding to terms such as “urban” and “suburban” were classified as urban areas, whereas those corresponding to terms such as “wild”, “nature reserve”, and “rural” were classified as natural areas. Classification of habitat types was confirmed using satellite imagery from Google Earth. Areas dominated by human infrastructure and artificial substrates were classified as urban areas, while those lacking such infrastructure and primarily consisting of forests, forest fragments, and agricultural land were classified as natural areas.

Geographic distribution data on our seven urban-adapted species were sourced from the Global Biodiversity Information Facility (https://www.gbif.org/)^35^, including records from smartphone photos, zoos, and animal resource centers. Data on viruses associated with these urban-adapted species were obtained from The Global Virome in One Network (VIRION), the Enhanced Infectious Diseases Database (EID2), and GenBank^36,37^. The geographic distribution and host–virus interaction data were supplemented by records obtained from the literature. The distributions of these species and their associated viruses were mapped using ArcGIS (v10.8). The full genomic sequences of viruses associated with our urban-adapted species were retrieved from GenBank for phylogenetic analyses. Data regarding the host, location, virus classification, and submission date for each sequence were extracted. Virus names from all sources were standardized according to the NCBI Taxonomy database. The relationship between viral host range and multi-organ distribution of viruses was assessed using Spearman correlation analysis.

Viral records were also collected for domestic animals and humans to investigate potential virus transmission with urban-adapted wildlife. Twenty-one representative domestic animal species were included in the analyses, covering traditional livestock and specialty farmed animals. Their Latin and common names are provided in Table S1. Data on viruses associated with domestic animals and humans were obtained from the VIRION and EID2 database, and viral sequences were downloaded from GenBank.

We used generalized additive models (GAMs) fit using the *mgcv* package in R to assess temporal changes in the number of virus-related publications and number of viral sequences for our seven urban-adapted species^38^. GAMs included either publication or sequence counts as a negative binomial response and a nonlinear term of year using thin plate splines with smoothing penalty, fit using restricted maximum likelihood. To identify periods of significant change in publication or sequencing intensity, we computed the first derivative of predicted curves and assessed whether the corresponding 95% confidence interval (CI) of this derivative overlapped with zero^39^.

### Meta-analysis of virus prevalence and seroprevalence

We analyzed virus prevalence and seroprevalence separately: prevalence was classified from confirmation of infection using nucleic acid detection, microscopy, and virus isolation and culture, while seroprevalence was classified from confirmation of current or recent infection via antibodies. Only publications that reported the number of tested and positive samples were included. The combined prevalence or seroprevalence and 95% confidence interval (CI) were calculated for each virus family and species for each individual urban-adapted species and across all seven urban-adapted species. The prevalence or seroprevalence was provided without a 95% CI when only one publication was available for a given virus. Prevalence or seroprevalence at the virus family level were estimated using two approaches. If the original publication specified the total number of positive samples, prevalence or seroprevalence were calculated by dividing the total number of positives by the number of samples tested. If we could not determine the total number of positive samples due to co-existence of multiple viral species, the highest prevalence or seroprevalence for a viral species was used to represent the entire family’s prevalence or seroprevalence. The *I²* statistic was used to quantify heterogeneity in prevalence and seroprevalence. If *I²* was less than 50%, a fixed-effects model was applied. Otherwise, a random-effects model was used, with an observation-level random effect nested in a study-level random effect. The meta-analysis was conducted using the *metafor* package in R, using the Freeman-Tukey double arcsine transformation for proportions and corresponding sampling variances.

### Ecological analysis

To analyze effects of habitat of urban-adapted species on virus outcomes, habitat was treated as a binary variable (i.e., natural or urban). Differences in viral richness across habitats were analyzed using Fisher’s exact test. The relationship between viral richness and number of relevant publications was assessed using Spearman correlation analysis. Diagnostic methods (prevalence assays versus seroprevalence assays) weakly affected virus positivity in these seven urban-adapted species (multilevel mixed linear model: total viruses, β = 0.026, *P* = 0.10; human-associated viruses: β = 0.067, *P* = 0.18). Therefore, prevalence (102 records) and seroprevalence (182 records) data were combined to analyze the impact of urbanization on virus positivity. After excluding positivity records with fewer than ten samples tested, five urban-adapted species (raccoon dog, raccoon, red fox, wild boar, and European shrew) were retained for analysis. The European shrew was further removed due to virus positivity in urban areas being consistently zero. To account for the potential interaction between habitat type and host species on positivity for total and human-associated viruses, generalized linear mixed models (GLMMs) were applied to fit two models, one with and one without an interaction term; these two models were compared with a likelihood ratio test. Parameters were estimated using maximum likelihood. The binomial response variables were positivity of total and human-associated viruses, respectively. Fixed effects included habitat type and host species, while virus family was treated as a random intercept to reflect the potential impact of different viral families on virus positivity. The overall statistic for the interaction term was estimated using *car* package in R, while the estimated slopes and multiplicity-adjusted *P*-values for each species in urban areas were obtained using *emmeans* package in R.

### Phylogenetic analyses

Phylogenetic analyses were performed for human-associated viruses harbored by the urban-adapted mammal species. Human-associated viruses were defined as those originating from humans in the VIRION, EID2, or GenBank databases. The coding sequences of the RNA-directed RNA polymerase (RdRp) gene and DNA polymerase (DPOL) genes were used for RNA viruses and DNA viruses, respectively. For viruses lacking these genes, other specific genes were used, such as the hemagglutinin (HA) gene for influenza viruses and the spike (S) gene for coronaviruses (see Table S2 for a full list of genes used for each virus). Sequence alignments were performed using MAFFT (v7.505)^40^. Ambiguous regions of each alignment were trimmed using BioEdit (v7.7.1.0). Maximum likelihood phylogenetic trees were generated using IQ-TREE (v2.1.4)^41^, implementing the best-fit nucleotide substitution model determined by the Bayesian information criterion score. Branch support was evaluated using 1,000 SH-like approximate likelihood ratio test (SH-aLRT) replicates. Phylogenetic trees were visualized using *ggtree* (v3.12.0) and *ggplot2* (v3.5.1) packages in R.

### Virus-sharing network

Virus-sharing networks were constructed using Cytoscape (v3.9.1)^42^. An edge connected two host species if they shared at least one virus. Edge thickness represented the number of shared viruses between two species. Our seven urban-adapted species as well as humans and the 21 domestic animal species were included in the analysis. Eigenvector centrality was used to assess the centrality of each host species, with centrality weighted by the number of viruses shared among host species. Domestic animals were either treated as separate nodes or grouped as a single entity for constructing the network.

## Results

### Description of the urban-adapted species dataset

We collected data from multiple sources and compiled a dataset of records for seven priority urban-adapted mammal species and their associated viruses from over 115 countries globally spanning 1574 to 2023. The dataset included 1,625,273 geographic location records of these urban-adapted species, 18,668 records of their associated viruses, and 14,356 viral sequences. We observed a significant non-linear increase in the number of virus-related publications (χ^2^_3.92,9_ = 758.8, *p* < 0.001) and viral sequences (χ^2^_1.42,9_ = 139.4, *p* < 0.001), with the greatest periods of change beginning in the 1950s and early 2000s, respectively; both measures of research intensity continued to significantly increase after the two coronavirus pandemics in 2003 and 2019 (Fig. 1a).

**Figure 1.**
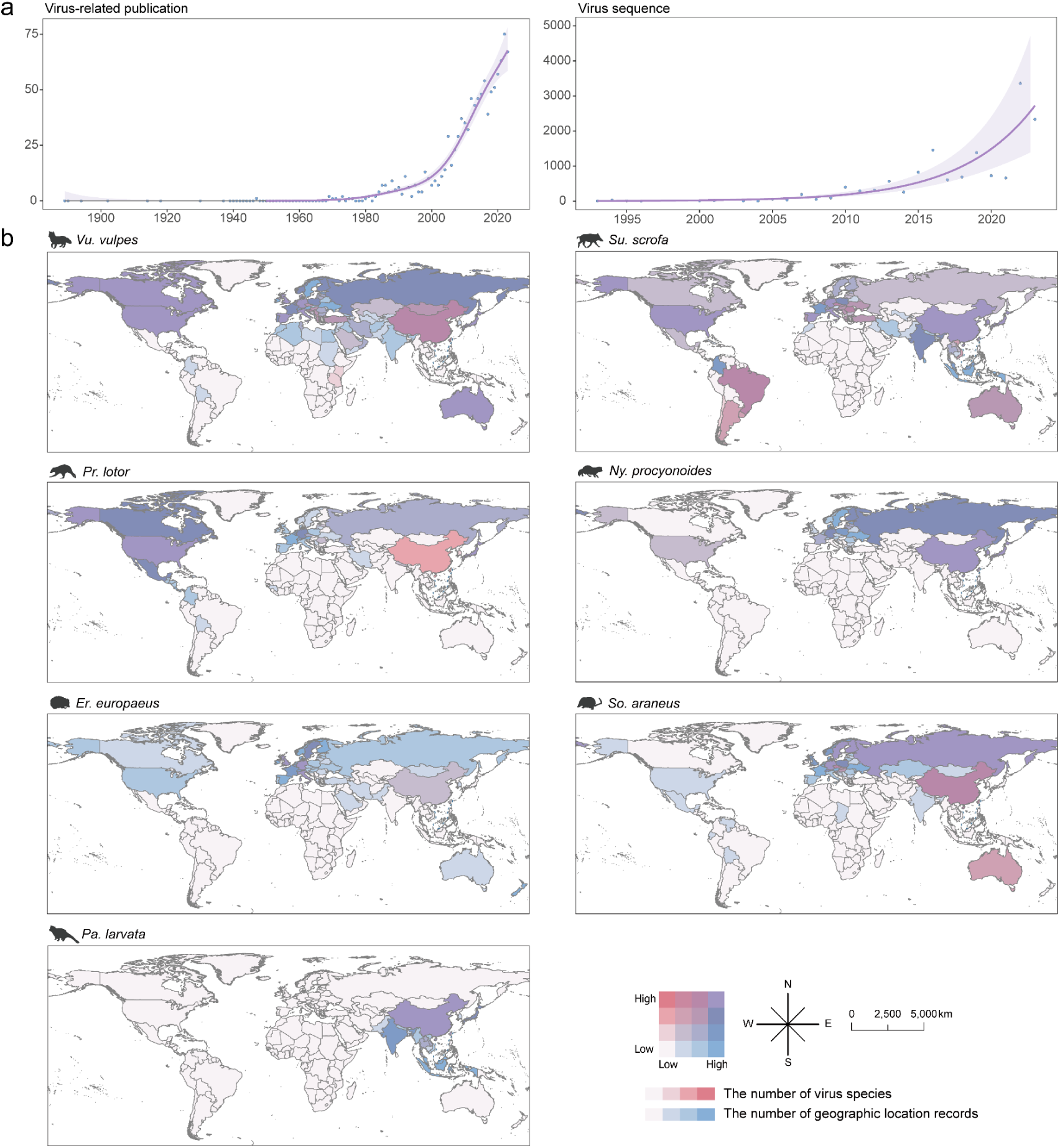
Temporal trends and geographic distribution of seven priority urban-adapted species and their associated viruses. a) Trends in the number of virus-related publications and viral sequences over time for urban-adapted species. Points show the number of virus-related publications and virus sequences for urban-adapted species per year. Lines and shading show the fitted GAM trend (mean and pointwise 95% CI). Line color indicates periods with strong evidence of either an upward (purple) trend in growth rates (95% CI of the first derivative of the fitted GAM trend above zero) or no significant trend (grey). b) Geographical distribution of urban-adapted species and their associated viruses. Darker shades indicate areas with a higher number of recorded urban-adapted species and their associated viruses.

Overall, the seven urban-adapted species were recorded globally, although their distribution patterns varied (Fig. 1b and S2a). Red foxes and wild boars were more widely distributed than other species, appearing in Europe, Oceania, North America, Asia, Africa, and South America. Raccoons had abundant records in their native range (i.e., the United States and Canada) and were later introduced into European countries, such as Belgium and Germany. The raccoon dog was most commonly recorded in East Asia, with high frequencies in South Korea and Japan. The European hedgehog and European shrew were primarily recorded across Europe, while palm civets were documented in Japan, China, and neighboring countries.

### Viral diversity and prevalence/seroprevalence

We found a diverse array of viruses present in the seven urban-adapted species, with 286 virus species across 24 orders and 38 families (Fig. 2a). Red foxes (n = 86 viruses) carried the largest number of viruses, followed by wild boars (n = 78), raccoons (n = 76), raccoon dogs (n = 55), European shrews (n = 42), palm civets (n = 31), and European hedgehogs (n = 22; Fig. 2b). Urban-adapted species within the same order tended to share similar virus compositions (Fig. 2c). Viruses from the families *Parvoviridae*, *Picornaviridae*, *Coronaviridae*, *Astroviridae*, *Flaviviridae*, *Circoviridae*, *Orthoherpesviridae*, and *Rhabdoviridae* were most commonly detected (i.e., in more than four of the urban-adapted species), accounting for 50% of all viruses (Fig. 2d). Other viruses were detected more sporadically; for example, viruses in the families *Togaviridae* and *Smacoviridae* were mainly identified in the raccoon and raccoon dog, respectively. The geographical distribution of viruses was generally consistent with that of the urban-adapted species, with the exception of raccoons (Fig. S2a-b). Raccoons were mainly recorded in North America and Europe but displayed the highest viral richness in North America and Asia (Fig. 1b).

**Figure 2.**
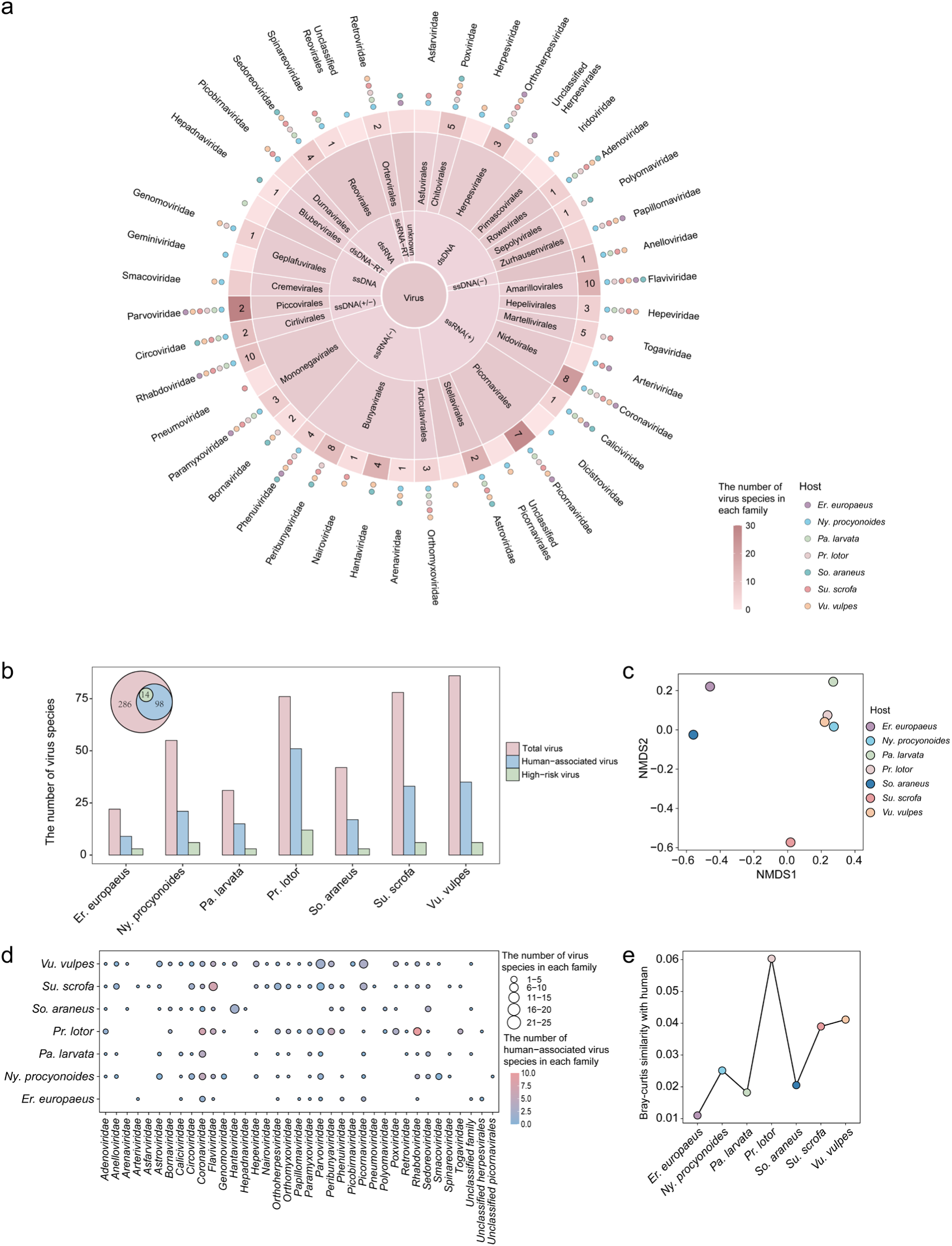
Overview of viruses harbored by urban-adapted species. a) Classification of viruses in urban-adapted species. The second and third inner circles represent the molecular type and order of viruses, respectively. In the grid of the fourth circle, gradient color represents the number of viruses within each family, and numbers represent the number of human-associated viruses within each family. The points outside the fourth circle represent urban-adapted species. The outermost names represent virus families. b) Comparison of number of total viruses, human-associated viruses, and high-risk viruses harbored by each urban-adapted species. c) Non-metric multidimensional scaling analysis showing the variation in virus composition at the species level among seven urban-adapted species. d) The number of viruses and human-associated viruses within each family in urban-adapted species. Circle size indicates the number of viruses, and circle color represents the number of human-associated viruses. e) The Bray-Curtis similarity of virus communities between each urban-adapted species and humans.

Several virus families showed high prevalence and seroprevalence in multiple urban-adapted species (Fig. 3a). Viruses in the family *Paramyxoviridae* were detected with high prevalence in palm civets (95%; 95% CI: 63–100%), raccoon dogs (52%; 95% CI: 28–75%), raccoons (50%; 95% CI: 34–66%), red foxes (30%; 95% CI: 14–47%), wild boars (21%; 95% CI: 13–30%), and European hedgehogs (9%; 95% CI: 0–42%). Viruses in the family *Rhabdoviridae*, primarily rabies virus, were found among hosts widely distributed across five continents, including raccoons (84% prevalence; 95% CI: 26–100%) and red foxes (31% seroprevalence; 95% CI: 8–60%; Fig. 3a and S3). In contrast, certain virus families were more specialized, with variable prevalence among virus species. For example, viruses in the family *Hantaviridae* were mainly detected in the European shrew (13%; 95% CI: 6–21%), with virus-specific prevalences of 25% (Asikkala virus), 16% (Seewis virus; 95% CI: 12–20%), and 2% (Seoul virus; 95% CI: 0–29%; Fig. S4).

**Figure 3.**
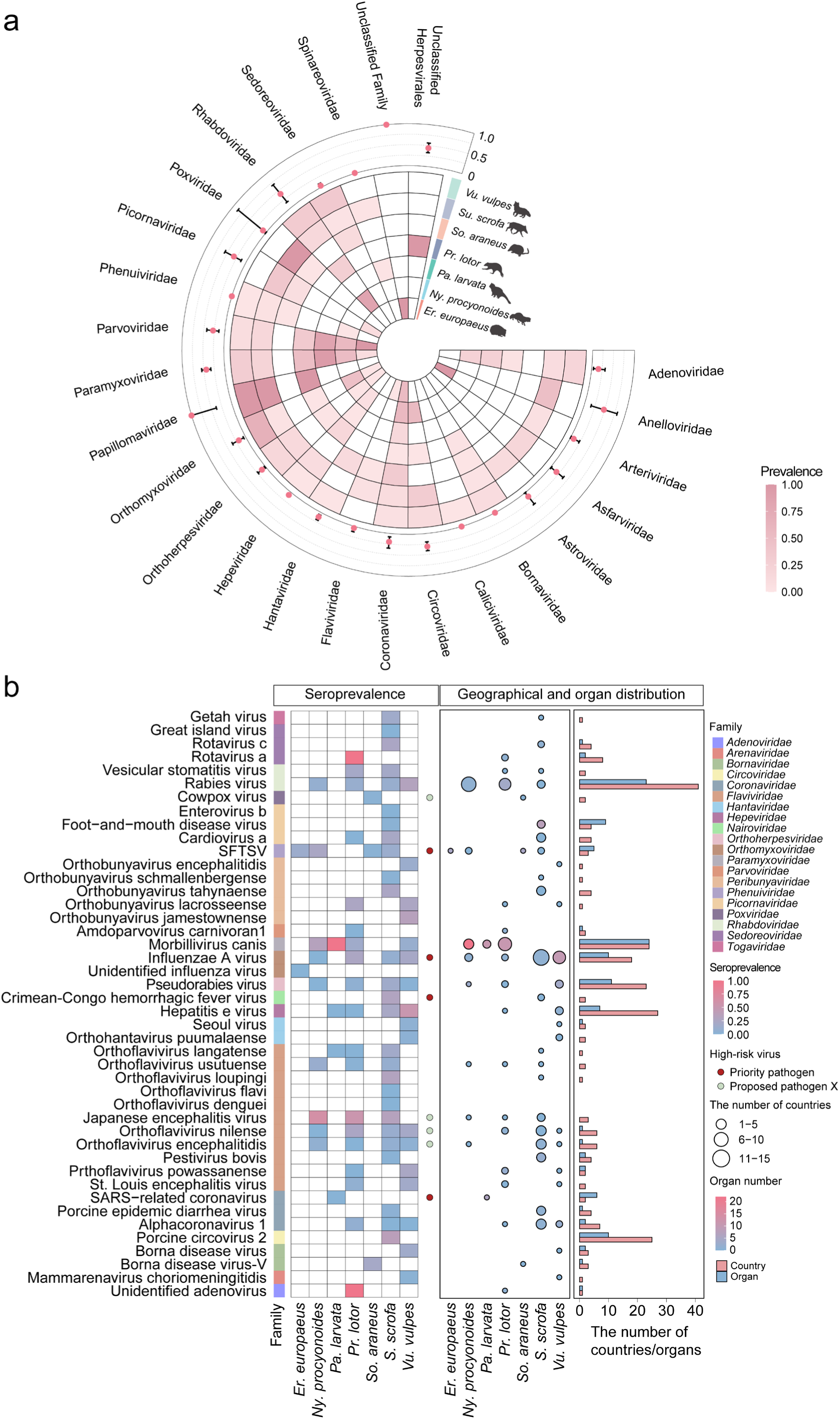
Positivity of viruses in urban-adapted species. a) Prevalence of viral families. The first to seventh inner circles represent the prevalence of each virus family in urban-adapted species. The outermost circle displays the combined prevalence and 95% CI for each virus family from meta-analysis. b) The epidemiological patterns of human-associated viruses. The first column lists the virus families. The heatmap displays the seroprevalence of human-associated viruses in each urban-adapted species, with points on the right side of the heatmap indicating high-risk viruses. The bubble chart shows the geographical and organ distribution of human-associated viruses in each urban-adapted species. Circle size represents the number of countries, and circle color represents the number of organs. The bar chart shows the combined number of countries and organs for each virus.

### Urban-adapted species harbor viruses with zoonotic potential

Approximately one-third of the viruses found in urban-adapted species (34.3%, 98/286) were associated with humans (Fig. 2b). Notably, fourteen viruses were classified by the World Health Organization as priority pathogens or potential pathogen X^43^. These viruses included influenza A virus, severe acute respiratory syndrome-related coronavirus, and Middle East respiratory syndrome-related coronavirus, which are highly transmissible and virulent, posing high risks for future outbreaks and pandemics. The virus community in raccoons was the most similar to that of humans (Fig. 2b and 2e). Raccoons carried the largest number of high-risk viruses (n = 12), followed by raccoon dogs (n = 6), wild boars (n = 6), red foxes (n = 6), palm civets (n = 3), European hedgehogs (n = 3), and European shrews (n = 3; Fig. 2b).

Several potentially high-risk viruses exhibited high prevalence or seroprevalence across multiple urban-adapted species or had a wide geographical distribution (Fig. 3b, S4, and S5). For example, influenza A virus was highly prevalent in red foxes (67%; 95% CI: 31–96%) and raccoon dogs (47%; 95% CI: 2–96%) and present but less prevalent in wild boars (3%; 95% CI: 0–10%), across 18 countries. Japanese encephalitis virus showed high seroprevalence in raccoon dogs (63%), raccoons (53%), and wild boars (33%; 95% CI: 12–57%). In contrast, Crimean-Congo hemorrhagic fever virus had a smaller host range, primarily detected in wild boar (29% seroprevalence; 95% CI: 11– 52%) in Spain and Poland. Notably, the same virus showed distinct intra-host organ distribution in different urban-adapted species. For instance, severe acute respiratory syndrome (SARS)-related coronavirus was detected in more organs in palm civet (8) than raccoon (4) and red fox (1). Viruses with a wide host range were also detected in more organs compared to rarely detected viruses (*ρ* = 0.72, *P* < 0.05).

**Figure 4.**
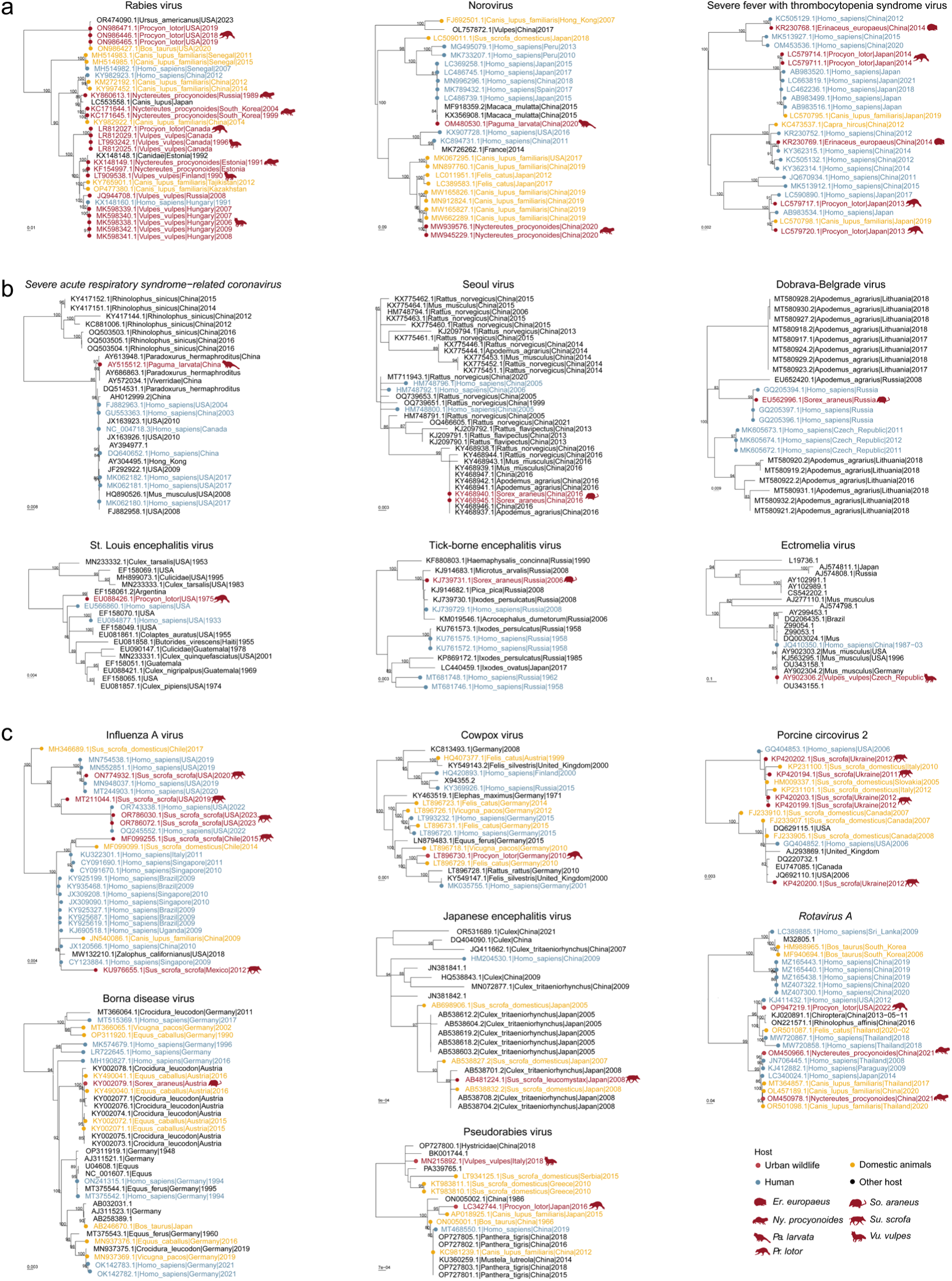
Phylogenetic trees of human-associated viruses in urban-adapted species. a) Potential bidirectional virus transmission between urban-adapted species and humans. b) Viruses from urban-adapted species cluster together with viruses found in humans. c) Viruses from urban-adapted species cluster together with viruses found in both humans and domestic animals. The trees were midpoint-rooted, with the scale bar representing the number of nucleotide substitutions per site. Only support values >80% are shown. Virus names include GenBank accession numbers, host species, sampling location, and sampling year.

### Urbanization affects virus diversity and positivity

We next investigated the effects of urbanization on viral richness and positivity in urban-adapted species. Viral richness was higher for hosts in urbanized areas than in natural areas (Fisher’s exact test: *P* = 0.004). In addition, viral richness across different habitat types correlated with the number of relevant publications (*ρ* = 0.919, *P* < 0.001). A likelihood ratio test showed significant differences for GLMMs with and without interaction terms (total viruses, χ^2^ = 253.26, *P* < 0.001; human-associated viruses, χ^2^ = 167.19, *P* < 0.001), suggesting interactions between host and habitat on virus positivity (Fig. 6a). Therefore, subsequent analyses were based on results from GLMMs with this interaction term.

Across total viruses, the effect of habitat type on virus positivity varied by host species (χ^2^ = 279.13, *P* < 0.001; Fig. 6b and Fig. 6c). Specifically, raccoon dogs (β = 1.28, *P* < 0.001) and red foxes (β = 0.75, *P* < 0.001) showed higher positivity in urbanized areas. In contrast, raccoons (β = -1.33, *P* < 0.001) and wild boar (β = -0.84, *P* < 0.001) exhibited lower positivity in urban compared to natural areas. Regarding human-associated viruses, the effect of habitat type on virus positivity also varied by host species (χ^2^ = 218.34, *P* < 0.001; Fig. 6b and Fig. 6c). Raccoon dogs (β = 1.92, *P* < 0.001) showed higher virus positivity in urban areas, whereas raccoons (β = -1.54, *P* < 0.001) and wild boar (β = -1.26, *P* < 0.001) had lower positivity in urban areas. No significant difference was observed between red foxes in natural and urban areas for human-associated viruses (β = 0.40, *P* = 0.098; Fig. 6b and Fig. 6c). Virus positivity also varied across different virus families (Fig. S6).

### Transmission and sharing of human-associated viruses

Phylogenetic trees revealed a close evolutionary relationship between human-associated viruses identified in urban-adapted species and humans. We observed multiple cases of potential bi-directional virus transmission between urban-adapted species and humans (Fig. 4a). Rabies viruses from red foxes and humans exhibited close phylogenetic relationships (RdRp gene: 99.4% nucleotide identity). Red foxes are known reservoir hosts for rabies virus and can transmit the virus to humans through bites or scratches^44^. Moreover, the GII.17 norovirus from palm civets in China was closely related to strains from humans (RdRp gene: 98.5–99.4% nucleotide identity); this likely indicates spillback, as the virus mainly circulates in humans^45^. SFTSV is an emerging tick-borne virus with a case-fatality rate up to 30%^46^. SFTSV from raccoons and European hedgehogs were grouped with human strains in distinct phylogenetic clusters (RdRp gene: 98.1–99.77% nucleotide identity; 99.4–100% nucleotide identity).

While *Haemaphysalis longicornis* is the primary vector of SFTSV, hedgehogs may serve as amplifying hosts^32^. Six human-associated viruses from urban-adapted species exhibited close evolutionary relationships with those found in humans (Fig. 4b). SARS-related coronaviruses in palm civets were genetically clustered in the same clade with human-origin strains (S gene: 99.1–99.3% nucleotide identity). Two vector-borne viruses, tick-borne encephalitis virus from European shrews and St. Louis encephalitis virus from raccoons, were clustered with human-origin viruses (RdRp gene: 99.6% nucleotide identity; 97.4–97.9% nucleotide identity). In addition, two hemorrhagic fever with renal syndrome viruses from European shrews, Dobrava-Belgrade virus and Seoul virus, exhibited close phylogenetic relationships with human viruses from the same country (nucleocapsid gene: 95.6–96.3% nucleotide identity; RdRp gene: 96.6–97.2% nucleotide identity). Ectromelia virus from red foxes was phylogenetically related to human-origin viruses (HA gene: 99.8% nucleotide identity).

Seven human-associated viruses from urban-adapted species clustered with viruses from both domestic animals and humans (Fig. 4c). Influenza A viruses from wild boars, domestic pigs, and humans exhibited close phylogenetic relationships (HA gene: 97– 98.4% nucleotide identity). Borna disease virus from European shrews was closely related to sequences from horses (*Equus caballus*) and humans (RdRp gene: 97.1–100% nucleotide identity). Porcine circovirus 2 from wild boars was clustered with sequences from domestic pigs and humans (DPOL gene: 99–99.5% nucleotide identity). Similar patterns were observed for rotavirus A, cowpox virus, pseudorabies virus, and the mosquito-borne Japanese encephalitis virus.

We further investigated the general pattern of virus-sharing among urban-adapted species, domestic animals, and humans. Analyses of the virus-sharing network revealed that domestic animals contribute heavily to virus transmission between urban-adapted wildlife and humans, with 82% of the viruses shared via domestic animals (Fig. 5a). Wild boars and raccoons shared more viruses with domestic animals than other urban-adapted species (Kruskal-Wallis test, *P* < 0.001; Fig. 5b). Among domestic animals, sheep (*Ovis aries*) and cows (*Bos taurus*) shared more viruses with urban-adapted species (Kruskal-Wallis test, *P* < 0.001; Fig. 5c and S7) and had higher weighted eigenvector centrality in the network. Viruses were less likely to be shared among the seven urban-adapted species studied, with only 40/98 (40.8%) viruses shared among them.

**Figure 5.**
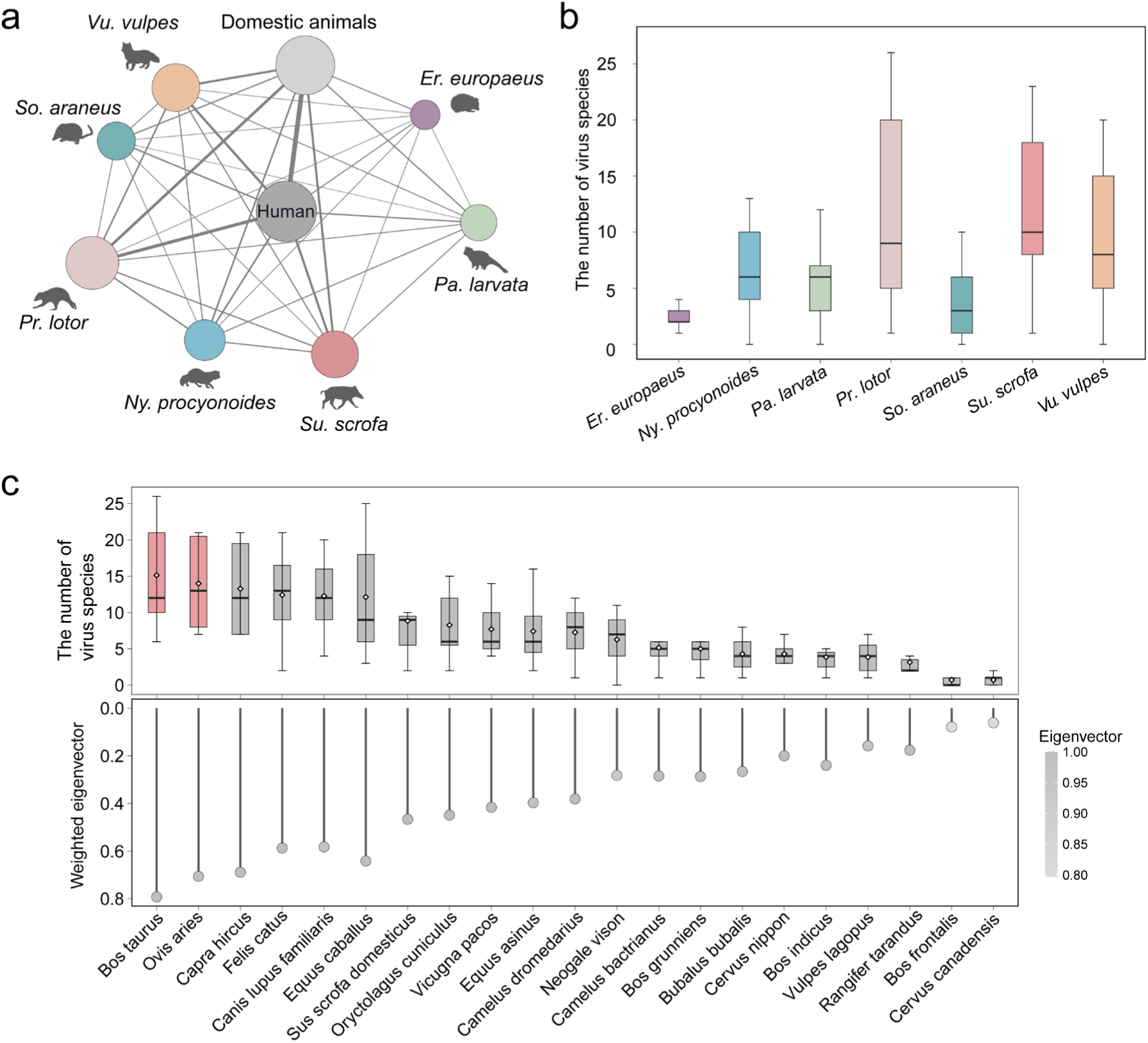
Virus transmission among urban-adapted species, domestic animals, and humans. a) The human-associated virus-sharing network. Node size indicates the number of human-associated viruses carried by each urban-adapted species. The edge thickness represents the number of shared viruses between two hosts. b) Boxplot showing the number of shared human-associated viruses between each urban-adapted species and domestic animals. c) Boxplot showing the number of shared viruses between each domestic animal and urban-adapted species. The horizontal line and rhombus represent the median and average number of shared viruses between each domestic animal and urban-adapted species, respectively. The lollipop chart displays the eigenvector centrality of each domestic animal species, with circle color representing the eigenvector centrality and the y-axis representing the weighted eigenvector centrality.

**Figure 6.**
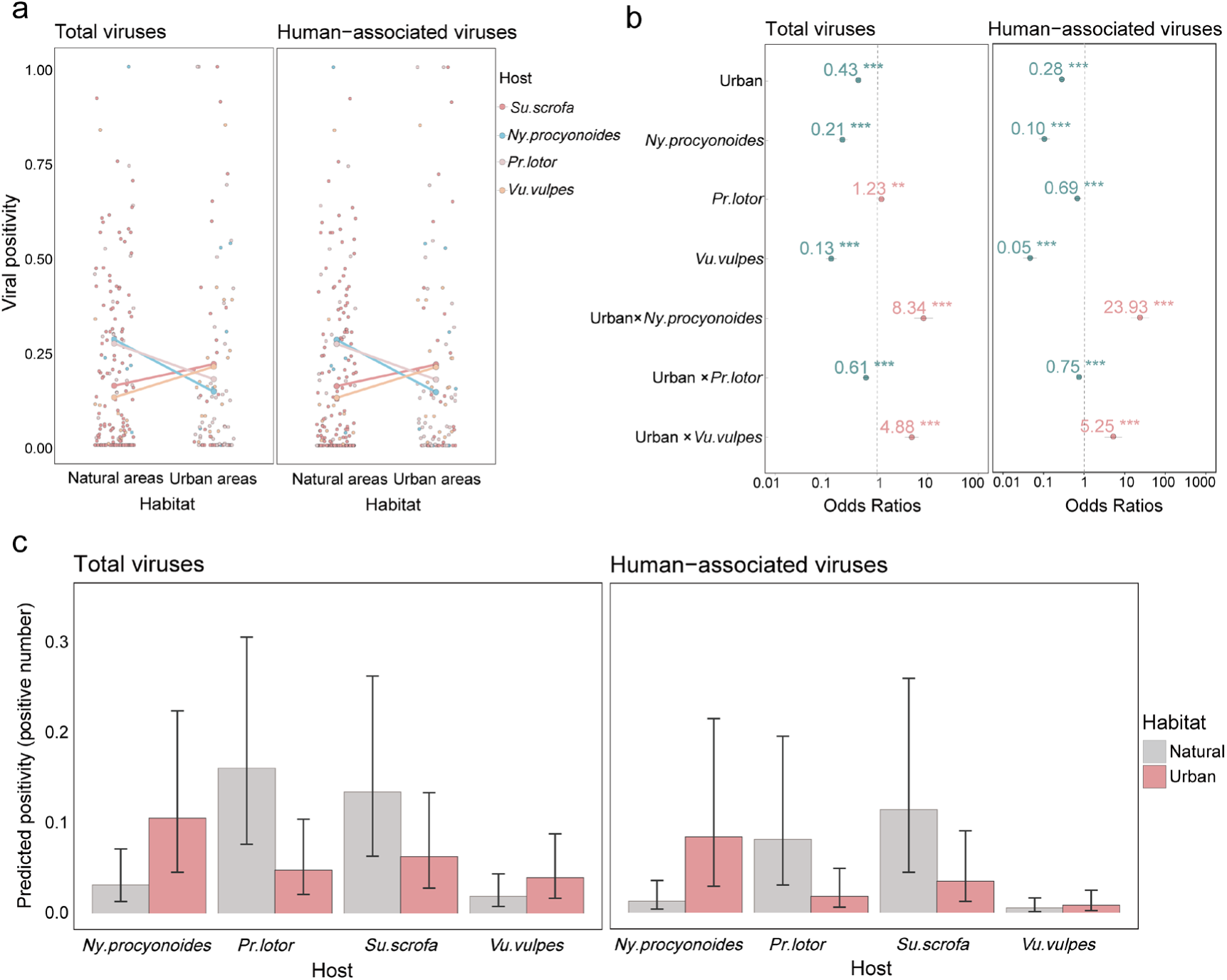
Virus positivity in urban and natural areas within urban-adapted species. a) Habitat-by-host interaction plots. b) Coefficients and corresponding *P*-values were estimated by the GLMMs for total and human-associated viruses. Coefficients are the odds ratios (OR) transformed by the exponential function (e). Red represents positive effects, while blue represents negative effects. Asterisks denote levels of statistical significance (***: *P*-values < 0.001, **: *P*-values < 0.01, *: *P*-values < 0.05). c) Predicted positivity for total and human-associated viruses from the GLMMs.

## Discussion

The repeated emergence of virulent and even pandemic viruses underscores the linkages between land-use change and wildlife disease dynamics, necessitating a deeper understanding of the role urban-adapted species play in viral transmission risks. In this study, we conducted an extensive investigation into the viruses of seven priority urban-adapted mammal species, encompassing viral diversity, viral prevalence and seroprevalence, effects of urbanization on viral positivity, and the risk of potential zoonotic spillover to and spillback from humans.

### Urban-adapted species harbor diverse zoonotic viruses

Our data showed that the seven urban-adapted species carried 98 human-associated viruses, of which 14 have been identified by the World Health Organization as high-risk viruses that could cause future outbreaks and pandemics. Urban-adapted species have been involved in the zoonotic transmission of multiple viruses, including hepatitis E virus, influenza A virus, and pseudorabies virus^47,48^. The synurbization of wildlife could increase their interactions with humans^7,8,49,50^. In urban settings, these species often exploit anthropogenic food resources, which can increase population size and density, promote host aggregation and intra-specific contact, and modify susceptibility to infection, all of which can amplify virus transmission^51,52^. We also found that these urban-adapted species share multiple viruses with domestic animals and companion animals. Interactions among urban-adapted species, domestic animals, and humans could thereby facilitate virus spillover to humans. Considering their rapid range expansion and population growth over the past decades^3^, urban-adapted species have the potential to act as unmonitored reservoirs for viruses of public health concern. Comprehensive surveillance that integrates urban-adapted species, domestic animals, and humans could inform risk mitigation strategies aimed at reducing zoonotic spillover events and protecting human health.

Influenza A viruses were prevalent in multiple urban-adapted species, including red foxes, raccoons, raccoon dogs, and wild boars. Urban-adapted species are free ranging and have numerous opportunities for exposure to influenza viruses, including avian influenza viruses. The HPAI H5N1 virus has been identified in red foxes^31^ and raccoon dogs^53^. In red foxes, the H5N1 virus has acquired the mammalian adaptation E627K in the polymerase basic two protein^19^. Moreover, urban-adapted species can be infected with both mammalian and avian influenza viruses, such as swine and avian influenza A viruses in wild boars^54^, and human and avian influenza A viruses in raccoons. Both the 1957 and 1968 influenza A virus pandemic strains likely arose through reassortment between an avian influenza strain and the strain circulating in humans at the time^55–57^. It is possible for urban-adapted species to serve as incubators facilitating the generation of genetically novel and potentially pandemic strains through reassortment, which could periodically spill over into humans, causing localized outbreaks or global pandemics.

### Virus spillback from urban-adapted species to humans

We found support for bidirectional virus transmission between urban-adapted wildlife and humans^58^. For example, we observed potential spillback of GII.17 norovirus from humans to palm civets. Spillback of virus from humans to wildlife can establish reservoirs with distinct evolutionary trajectories, ultimately impacting both animal and human health^15^. SARS-CoV-2 has been frequently transmitted from humans into wild deer, sustained efficient deer-to-deer transmission, and spilled back from wild deer to humans in the United States^14,59^. The virus evolved three times faster in the wild deer and acquired multiple adaptive mutations across the genome^13^. Urban-adapted wildlife could thus serve as breeding grounds for novel virus variants with unpredictable characteristics, posing continuous risks of zoonotic transmission back to humans.

### Virus positivity in urbanized and natural areas

We analyzed virus positivity proportions from four of our priority urban-adapted species to assess whether urban land use had consistent or species-specific effects on viral exposure or infection risk. For both all viruses and human-associated viruses, we found that urban impacts depended on the mammal species. Raccoon dogs had greater viral positivity in urban habitats, while both raccoons and wild boar had lower viral positivity in urban habitats; these effects were consistent among all viruses and only human-associated viruses. Red foxes also showed increased viral positivity in urban habitats, but only when considering all viruses (not the human-associated subset). Because sample size limitations necessitated pooling both prevalence and seroprevalence, such results may more likely indicate increases or decreases in viral exposure with urbanization. These findings provide further nuance to broader meta-analyses in which effects of urbanization vary across host groups; for example, such analyses have found that carnivores and primates have lower prevalence of complex life cycle parasites in urban habitats^26^. Yet other analyses have found that coarse taxonomic groupings have not explained variation in infection outcomes of urbanization^60^; such results could be partly explained by our findings here in which effects are highly species-specific. Greater attention to characteristics of species that facilitate increases or decreases in viral positivity with urbanization would provide more predictive insights^61^.

Despite uncertainty in the traits underlying these species-specific differences, our results also emphasize that urban populations of raccoon dogs could be an important target for viral surveillance and preempting spillover risks. The propensity of this species to capitalize on anthropogenic resources in urban habitats could increase their exposure to humans and domestic animals such as cats and dogs, providing conduits for cross-species transmission^62,63^. Further studies of viral dynamics in raccoon dogs, including longitudinal surveys of both natural and urbanized populations, could enhance understanding how urban habitats increase viral transmission^64^.

### Limitations

This study has several limitations that should be taken into consideration when interpreting the results. First, we include only literature published in English and Chinese. Second, current detection methods were not able to identify all potential pathogens in urban-adapted species. Pooled samples may reduce the sensitivity of virus discovery, and viruses of low abundance might go undetected. Finally, we selected seven priority urban-adapted mammal species based on their relatively large population size, wide geographic distribution, and previous association with zoonotic viruses. As these species were selected to be representative of urban-adapted mammals, we were not able to include all urban-adapted species in our study. Some other epidemiologically important and equally widespread urban-adapted species, such as coyotes (*Canis latrans*), Norway rats (*Rattus norvegicus*), and big brown bats (*Eptesicus fuscus*)^65–67^, require further investigation and would be good targets for further data syntheses.

## Conclusions

In summary, our findings demonstrate that urban-adapted wildlife carry a wide spectrum of zoonotic viruses, including some with a high risk of triggering future pandemics through viral spillover at the wildlife–human and wildlife–domestic animal– human interfaces. Enhanced surveillance of viruses circulating at these interfaces in urban settings will be crucial for providing early warnings about potential zoonotic spillover. Urban-adapted wildlife should thus be considered a critical component of public health interventions aimed at controlling emerging infectious diseases.

## Acknowledgements

This study was supported by the National Natural Science Foundation of China (32470561, 32271605), Taishan Scholars Project (tsqn202306003), and Shandong Excellent Young Scientists Fund Program (2022HWYQ-056). DJB was supported by the National Science Foundation (BII 2213854).

## Competing interests

None declared.

## Notes

### Competing Interest Statement

The authors have declared no competing interest.

## Reference

1. Magle, S. B., Hunt, V. M., Vernon, M. & Crooks, K. R. Urban wildlife research: Past, present, and future. Biol Conserv 155, 23–32 (2012).

2. Santini, L. et al. One strategy does not fit all: determinants of urban adaptation in mammals. Ecol Lett 22, 365–376 (2019).

3. Ma, D. et al. Global expansion of human-wildlife overlap in the 21st century. Sci Adv 10, eadp7706 (2024).

4. Jones, K. E. et al. Global trends in emerging infectious diseases. Nature 451, 990– 993 (2008).

5. Olival, K. J. et al. Host and viral traits predict zoonotic spillover from mammals. Nature 546, 646–650 (2017).

6. Plowright, R. K. et al. Land use-induced spillover: a call to action to safeguard environmental, animal, and human health. Lancet Planet Health 5, e237–e245 (2021).

7. Betke, B. A., Gottdenker, N. L., Meyers, L. A. & Becker, D. J. Ecological and evolutionary characteristics of anthropogenic roosting ability in bats of the world. iScience 27, 110369 (2024).

8. Ecke, F. et al. Population fluctuations and synanthropy explain transmission risk in rodent-borne zoonoses. Nat Commun 13, 7532 (2022).

9. Xiao, X., Newman, C., Buesching, C. D., Macdonald, D. W. & Zhou, Z.-M. Animal sales from Wuhan wet markets immediately prior to the COVID-19 pandemic. Sci Rep 11, 11898 (2021).

10. Zhao, J., Cui, W. & Tian, B. The Potential Intermediate Hosts for SARS-CoV-2. Front Microbiol 11, (2020).

11. Hassell, J. M., Begon, M., Ward, M. J. & Fèvre, E. M. Urbanization and Disease Emergence: Dynamics at the Wildlife–Livestock–Human Interface. Trends Ecol Evol 32, 55–67 (2017).

12. Blasdell, K. R. et al. Rats and the city: Implications of urbanization on zoonotic disease risk in Southeast Asia. Proceedings of the National Academy of Sciences 119, e2112341119 (2022).

13. McBride, D. S. et al. Accelerated evolution of SARS-CoV-2 in free-ranging white-tailed deer. Nat Commun 14, 5105 (2023).

14. Feng, A. et al. Transmission of SARS-CoV-2 in free-ranging white-tailed deer in the United States. Nat Commun 14, 4078 (2023).

15. Fagre, A. C. et al. Assessing the risk of human-to-wildlife pathogen transmission for conservation and public health. Ecol Lett 25, 1534–1549 (2022).

16. Rasche, A. et al. Highly diversified shrew hepatitis B viruses corroborate ancient origins and divergent infection patterns of mammalian hepadnaviruses. Proceedings of the National Academy of Sciences 116, 17007–17012 (2019).

17. Goethert, H. K., Mather, T. N., Johnson, R. W. & Telford, S. R. Incrimination of shrews as a reservoir for Powassan virus. Commun Biol 4, 1319 (2021).

18. Campbell, S. J. et al. Red fox viromes in urban and rural landscapes. Virus Evol 6, veaa065 (2020).

19. Luca, B. et al. Highly Pathogenic Avian Influenza H5N1 Virus Infections in Wild Red Foxes (Vulpes vulpes) Show Neurotropism and Adaptive Virus Mutations. Microbiol Spectr 11, e02867–22 (2023).

20. Fisher, A. M. et al. The ecology of viruses in urban rodents with a focus on SARS-CoV-2. Emerg Microbes Infect 12, 2217940 (2023).

21. Schotte, U. et al. Phylogeny and spatiotemporal dynamics of hepatitis E virus infections in wild boar and deer from six areas of Germany during 2013–2017. Transbound Emerg Dis 69, e1992–e2005 (2022).

22. Wei-Shan, C. et al. Metatranscriptomic Analysis of Virus Diversity in Urban Wild Birds with Paretic Disease. J Virol 94, 10.1128/jvi.00606-20 (2020).

23. Denstedt, E. et al. Detection of African swine fever virus in free-ranging wild boar in Southeast Asia. Transbound Emerg Dis 68, 2669–2675 (2021).

24. Song, T. et al. First detection and phylogenetic analysis of porcine circovirus type 2 in raccoon dogs. BMC Vet Res 15, 107 (2019).

25. Albery, G. F. et al. Urban-adapted mammal species have more known pathogens. Nat Ecol Evol 6, 794–801 (2022).

26. Werner, C. S. & Nunn, C. L. Effect of urban habitat use on parasitism in mammals: a meta-analysis. Proc Biol Sci 287, 20200397 (2020).

27. Snow, N. P., Jarzyna, M. A. & VerCauteren, K. C. Interpreting and predicting the spread of invasive wild pigs. Journal of Applied Ecology 54, 2022–2032 (2017).

28. Jackowiak, M. et al. Colonization of Warsaw by the red fox Vulpes vulpes in the years 1976-2019. Sci Rep 11, 13931 (2021).

29. Fischer, M. L. et al. Assessing and predicting the spread of non-native raccoons in Germany using hunting bag data and dispersal weighted models. Biol Invasions 18, 57–71 (2016).

30. Myśliwy, I., Perec-Matysiak, A. & Hildebrand, J. Invasive raccoon (Procyon lotor) and raccoon dog (Nyctereutes procyonoides) as potential reservoirs of tick-borne pathogens: data review from native and introduced areas. Parasit Vectors 15, 126 (2022).

31. Rijks, J. M. et al. Highly Pathogenic Avian Influenza A(H5N1) Virus in Wild Red Foxes, the Netherlands, 2021. Emerg Infect Dis **27**, 2960–2962 (2021).

32. Zhao, C. et al. Hedgehogs as Amplifying Hosts of Severe Fever with Thrombocytopenia Syndrome Virus, China. Emerging Infectious Disease journal 28, 2491 (2022).

33. Goldberg, A. R. et al. Widespread exposure to SARS-CoV-2 in wildlife communities. Nat Commun 15, 6210 (2024).

34. Page, M. J. et al. The PRISMA 2020 statement: an updated guideline for reporting systematic reviews. BMJ 372, n71 (2021).

35. GBIF.org. GBIF Occurrence Download 10.15468/dl.wr46fe. (2024) doi:10.1016/j.physbeh.2017.03.040.

36. Wardeh, M., Risley, C., McIntyre, M. K., Setzkorn, C. & Baylis, M. Database of host-pathogen and related species interactions, and their global distribution. Sci Data 2, 150049 (2015).

37. Carlson, C. J. et al. The Global Virome in One Network (VIRION): an Atlas of Vertebrate-Virus Associations. mBio 13, (2022).

38. Wood SN. Generalized additive models: an introduction with R. chapman and hall/CRC (2017).

39. Simpson, G. L. Modelling Palaeoecological Time Series Using Generalised Additive Models. Front Ecol Evol 6, (2018).

40. Katoh, K. & Standley, D. M. MAFFT Multiple Sequence Alignment Software Version 7: Improvements in Performance and Usability. Mol Biol Evol 30, 772–780 (2013).

41. Nguyen, L.-T., Schmidt, H. A., von Haeseler, A. & Minh, B. Q. IQ-TREE: A Fast and Effective Stochastic Algorithm for Estimating Maximum-Likelihood Phylogenies. Mol Biol Evol 32, 268–274 (2015).

42. Shannon, P. et al. Cytoscape: A software Environment for integrated models of biomolecular interaction networks. Genome Res 13, 2498–2504 (2003).

43. World Health Organization. Pathogens prioritization: a scientific framework for epidemic and pandemic research preparedness. https://www.who.int/publications/m/item/pathogens-prioritization-a-scientific-framework-for-epidemic-and-pandemic-research-preparedness. *World Health Organization* (2024).

44. Pastoret, P. P. & Brochier, B. Epidemiology and control of fox rabies in Europe. Vaccine 17, 1750–1754 (1999).

45. Jin, M. et al. Norovirus Outbreak Surveillance, China, 2016–2018. Emerging Infectious Disease journal 26, 437 (2020).

46. Xue-Jie, Y. et al. Fever with Thrombocytopenia Associated with a Novel Bunyavirus in China. New England Journal of Medicine 364, 1523–1532 (2024).

47. Fujimoto, Y., Inoue, H., Ozawa, M. & Matsuu, A. Serological survey of influenza A virus infection in Japanese wild boars (Sus scrofa leucomystax). Microbiol Immunol 63, 517–522 (2019).

48. Gong, W. et al. Genetic characterization of hepatitis E virus from wild boar in China. Transbound Emerg Dis 69, e3357–e3362 (2022).

49. Jackson, R. T., Webala, P. W., Ogola, J. G., Lunn, T. J. & Forbes, K. M. Roost selection by synanthropic bats in rural Kenya: implications for human–wildlife conflict and zoonotic pathogen spillover. R Soc Open Sci 10, 230578 (2023).

50. Balasubramaniam, K. N. et al. Impact of joint interactions with humans and social interactions with conspecifics on the risk of zooanthroponotic outbreaks among wildlife populations. Sci Rep 12, 11600 (2022).

51. Becker, D. J., Streicker, D. G. & Altizer, S. Linking anthropogenic resources to wildlife–pathogen dynamics: a review and meta-analysis. Ecol Lett 18, 483–495 (2015).

52. Altizer, S. et al. Food for contagion: synthesis and future directions for studying host–parasite responses to resource shifts in anthropogenic environments. Philosophical Transactions of the Royal Society B: Biological Sciences 373, 20170102 (2018).

53. Qi, X. et al. Molecular Characterization of Highly Pathogenic H5N1 Avian Influenza A Viruses Isolated from Raccoon Dogs in China. PLoS One 4, e4682- (2009).

54. E, M. B., et al. Feral Swine in the United States Have Been Exposed to both Avian and Swine Influenza A Viruses. Appl Environ Microbiol 83, e01346–17 (2017).

55. Schäffr, J. R. et al. Origin of the Pandemic 1957 H2 Influenza A Virus and the Persistence of Its Possible Progenitors in the Avian Reservoir. Virology 194, 781– 788 (1993).

56. Kawaoka, Y., Krauss, S. & Webster, R. G. Avian-to-human transmission of the PB1 gene of influenza A viruses in the 1957 and 1968 pandemics. J Virol 63, 4603–4608 (1989).

57. Bean, W. J. et al. Evolution of the H3 influenza virus hemagglutinin from human and nonhuman hosts. J Virol 66, 1129–1138 (1992).

58. Olival, K. J. et al. Possibility for reverse zoonotic transmission of SARS-CoV-2 to free-ranging wildlife: A case study of bats. PLoS Pathog 16, e1008758- (2020).

59. Pickering, B. et al. Divergent SARS-CoV-2 variant emerges in white-tailed deer with deer-to-human transmission. Nat Microbiol 7, 2011–2024 (2022).

60. Murray, M. H. et al. City sicker? A meta-analysis of wildlife health and urbanization. Front Ecol Environ 17, 575–583 (2019).

61. Becker, D. J., Streicker, D. G. & Altizer, S. Using host species traits to understand the consequences of resource provisioning for host-parasite interactions. J Anim Ecol 87, 511–525 (2018).

62. Wang, Y. et al. Behavioral plasticity of raccoon dogs (Nyctereutes procyonoides) provides new insights for urban wildlife management in metropolis Shanghai, China. Environmental Research Letters 19, 104063 (2024).

63. Süld, K. et al. An invasive vector of zoonotic disease sustained by anthropogenic resources: the raccoon dog in northern Europe. PLoS One 9, e96358 (2014).

64. Plowright, R. K., Becker, D. J., McCallum, H. & Manlove, K. R. Sampling to elucidate the dynamics of infections in reservoir hosts. Philos Trans R Soc Lond B Biol Sci 374, 20180336 (2019).

65. Malmlov, A., Breck, S., Fry, T. & Duncan, C. Serologic survey for cross-species pathogens in urban coyotes (Canis latrans), Colorado, USA. J Wildl Dis 50, 946–50 (2014).

66. Streicker, D. G., Franka, R., Jackson, F. R. & Rupprecht, C. E. Anthropogenic roost switching and rabies virus dynamics in house-roosting big brown bats. Vector Borne Zoonotic Dis 13, 498–504 (2013).

67. Firth, C. et al. Detection of zoonotic pathogens and characterization of novel viruses carried by commensal Rattus norvegicus in New York City. mBio 5, e01933–14 (2014).

